# Cryptic diversification proceeds despite historical and contemporary hybridization in Patagonian ants

**DOI:** 10.64898/2026.08.08.743675

**Authors:** Melisa Olave, Pablo Pessacq, Jérémy Gauthier, Fabiana Cuezzo, Julia Bilat, Danielle Anjos-Santos, Marcelo Pereda Gomez, Mariana Morando, Luciano J. Avila, Nadir Alvarez

**Author notes:** **Data accessibility** Genomic data is publicly available at NCBI Project XXX [to be included upon ms acceptance]. All datasets are available at Dryad XXXX [to be included upon ms acceptance]. **Code availability** All custom code developed for this article is freely available at www.github.com/melisaolave/mafalda. **Authors contributions** M.O., N.A. conceived the study and obtained funding. M.O. analyzed the data and drafted the manuscript. F.C. identified species. P.P., D.A.S., M.G.P. and L.J.A. performed field work. M.M and P.P. obtained funding for field trips. J.B. performed the lab work. N.A. and J.G. assisted with recommendations in analyses. All authors helped with comments and editions on the manuscript and read and approved the final manuscript. **Competing interests** Authors declare no competing interests.

## Abstract

Understanding how independently evolving lineages arise despite ongoing gene flow remains a central question in evolutionary biology. Although genomic studies increasingly suggest that hybridization can accompany diversification, empirical evidence from ecologically dominant insect groups remains limited. Here, we present the first population-scale phylogenomic analysis of Patagonian ants, sampling *Dorymyrmex* across approximately 450,000 km^2^. Using genome-wide SNPs, coalescent phylogenetics, phylogenetic networks, demographic modelling, species delimitation, and genome scans, we reconstruct the evolutionary history of this widespread genus. We discovered extensive cryptic diversity with strong genomic differentiation and detected both recent and historical hybridization, demonstrating that substantial genomic divergence accumulated despite recurrent gene flow events. Genome scans further identify candidate loci associated with adaptation to Patagonia’s contrasting environments, suggesting that ecological divergence contributed to lineage diversification. Our results show that cryptic diversification can proceed despite recurrent gene flow, supporting hybridization as an integral component of the diversification process. More broadly, this study illustrates how genome-scale data can reveal hidden biodiversity and the evolutionary processes shaping it in ecologically important but genomically understudied taxa.

## Main text

Understanding how independently evolving lineages arise despite recurrent gene flow remains a central challenge in evolutionary biology (Long, et al., 2026). Resolving these processes requires genome-scale datasets capable of distinguishing population structure, hybridization and independently evolving lineages, yet such datasets remain scarce for much of Earth’s biodiversity (Hohenlohe, et al., 2021). This gap is especially pronounced in the Global South, which holds the vast majority of the world’s species richness yet faces a critical deficit in advanced genomics research (Turchetto-Zolet, et al., 2013; Linck & Cadena, 2024). As a consequence, a large proportion of species on Earth remain undescribed (Li & Wiens, 2023), while traditional morphological approaches often fail to recognize independently evolving lineages masked as cryptic diversity (Cheng, et al., 2025). In this context, high-throughput genomics has emerged as an indispensable tool, offering the resolution necessary to delineate cryptic species boundaries, and accelerate biodiversity discovery before these undocumented lineages are lost to rapid environmental change (Costello, 2013).

Ants (Hymenoptera: Formicidae) are the world’s most successful eusocial insects. Having originated 99 million years ago in the Cretaceous from a wasp-like ancestor (Barden, 2017), ants have since diversified into the currently, there are 14,452 described species recognized today (AntCat, 2026), spreading across most terrestrial ecosystems, where they fill different ecological roles. Ants are estimated to have a biomass that exceeds the combined biomass of all wild birds and mammals on Earth (Schultheiss, et al., 2022). They dominate plant resource consumption in the canopies of lowland rainforests (Davidson, et al., 2003) and are ecosystem engineers, shaping community structure in innumerable ways. Ants negatively impact other animal species on which they prey or which they outcompete; create enemy free-space around animal species that they farm; act as hosts to animal social parasites; control bulk energy flows through defoliation, decomposition and bioturbation (influencing plant life and soil properties); impact seed dispersal (often positively through myrmecochory) and pollination (generally negatively through nectar theft and antimicrobial secretions that reduce pollen viability); and serve as a major food source for still other animals (Hölldobler & Wilson, 1990; Folgarait, 1998; Lach, et al., 2009; Elizalde, et al., 2020; Schultheiss, et al., 2022). Despite their ecological dominance, ants remain underrepresented in phylogenomic studies of diversification, limiting our understanding of how speciation and hybridization have shaped one of the most successful insect radiations

*Dorymyrmex* (Mayr, 1866) are ants characterized by monomorphic workers (61 valid species and 26 subspecies; (AntCat, 2026)) with very few diagnostic features, some of which are cryptic and not immediately evident through standard morphology (Oberski, 2022). This structural homogeneity, coupled with the vast, under-sampled territory of Patagonia and a shortage of local trained taxonomists, severely limits our knowledge of Patagonian *Dorymyrmex*. Originating 23 million years ago, this ant genus favor deserts, road-sides, and open grasslands, and are frequently encountered in open habitats across the Americas, from the Great Planes to the southernmost of Patagonia (Oberski, 2022; Oberski, 2024). *Dorymyrmex* ants are conspicuous in open habitats and build ground nests usually marked by craters or cones of soil, predominantly forage during daylight hours, and actively scavenge (Hölldobler & Wilson, 1990). Among ant species that have conquered the southernmost tip of the world, *Dorymyrmex tener* (Mayr, 1866), *Dorymyrmex richteri* (Forel, 1911) and *Dorymyrmex flavescens* (Mayr, 1866) are abundant across the Patagonian steppe. Despite their ecological importance, *Dorymyrmex* remains biologically known and (like much of the Global South’s insect fauna) the extent of its genetic diversity has never been examined with densely sampled genomic data. The limited morphological differentiation among *Dorymyrmex* species makes this genus particularly suitable for investigating how genomic divergence accumulates during diversification, including the extent to which cryptic lineages remain connected by gene flow.

Patagonia, a region of the South American Transition Zone (Roig-Juñent, et al., 2018), represents a natural laboratory for studying diversification and speciation dynamics due to the dramatic landscape alterations caused by glaciations, climate change, tectonics, volcanism, palaeobasins, and seashore shifts with marine transgressions (Ramos & Ghiglione, 2008; Martinez & Kutscher, 2011; Ponce, et al., 2011). Numerous glacial advances and retreats during the Pleistocene, such as the Greatest Patagonian Glaciation (GPG; 1.2 to 1.0 Mya) and the Last Glacial Maximum (LGM; 20 to 18 kya), have differentially impacted the local biota (McCulloch, et al., 2000; Coronato, et al., 2008; Ponce, et al., 2011). Yet, Patagonia remains as a very poorly known from a genomics standpoint. In addition, little genomic research has been conducted on Patagonian insects (beetles: (Olave, et al., 2023), grasshoppers: (Guzman, et al., 2023)), and only one study has addressed the genetics of ants (*Acromyrmex* (Mayr, 1866), (Sanchez-Restrepo, et al., 2023)), but based on one mitochondrial and four nuclear genes.

Here, we use genome-wide SNP data to investigate how genomic divergence accumulated during the diversification of Patagonian *Dorymyrmex* ants. Specifically, we test whether independently evolving lineages can arise despite recent and historical hybridization, reconstruct patterns of diversification across space and time, evaluate species boundaries using coalescent-based methods, and identify candidate loci associated with local adaptation. Because *Dorymyrmex* has never been examined using dense genomic sampling across Patagonia (∼450,000 km^2^), our study also provides the first comprehensive assessment of hidden evolutionary diversity in this ecologically important genus.

## Material and Methods

### Field sampling

Specimens of D. *tener, D. richteri and D. flavescens*, plus representatives of *Dorymyrmex sp*. and *Dorymyrmex pyramicus* (Roger, 1863) used as outgroups, were collected by hand or pitfall traps and stored in a freezer with 96% ethanol at the National University of Patagonia (UNPSJB, Argentina) Entomological collection. Sampling locations represent a wide range covering ∼450.000 km^2^ (linear range >2 500 km), from Mendoza to Santa Cruz provinces (Fig. 1a).

**Fig. 1:**
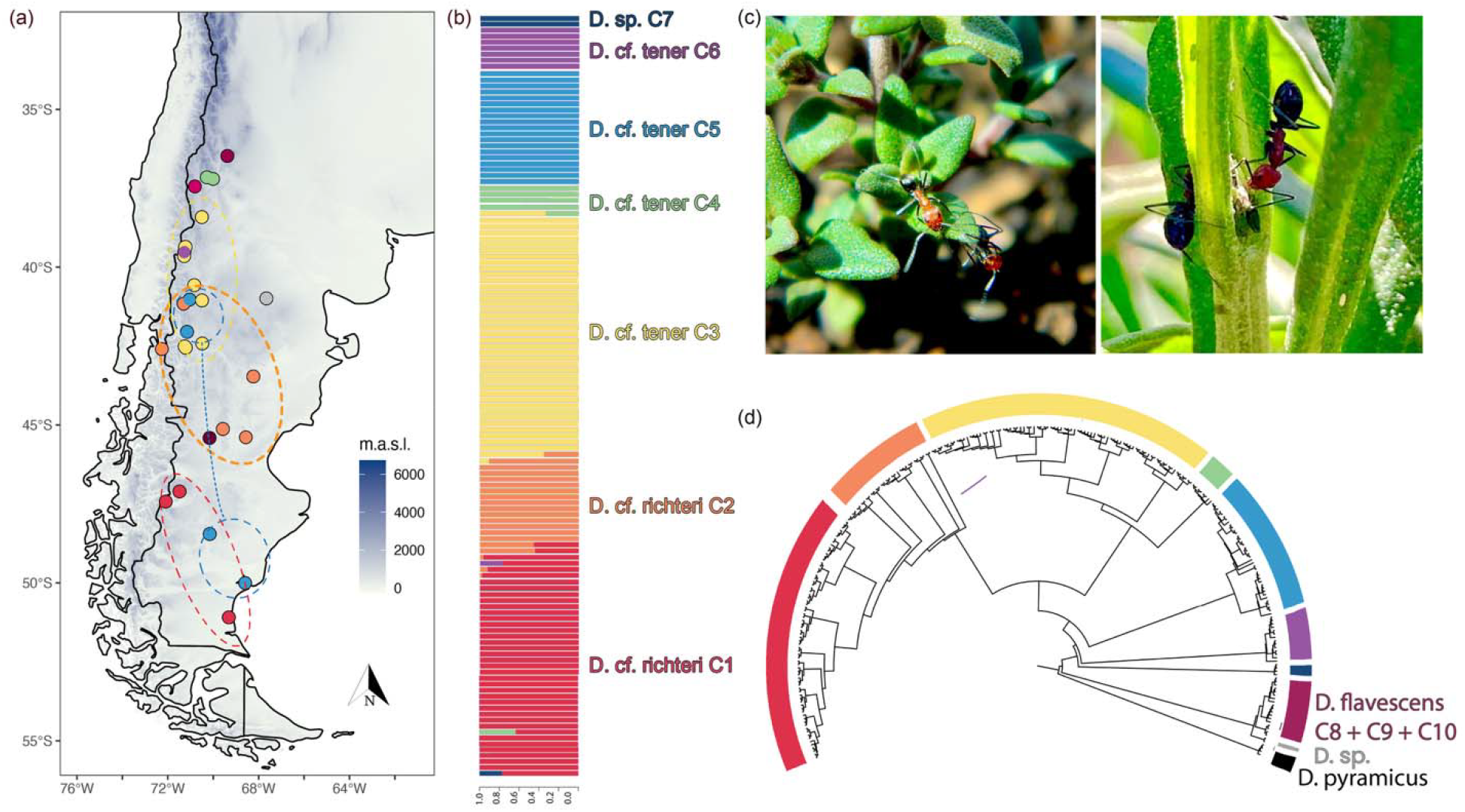
Sampling and exploratory analyses of genomic data. (a) sampling localities with color-code following clustering in (b) admixture analysis and also in (d) the neighbor joining (NJ) tree shown. Admixture plot constructed on 121,703 SNPs including 126 individuals. The NJ tree was obtained based on 19,121 SNPs for 276 alleles (138 diploid individuals). (c) Workers of *D.tener* in their natural environment (Credits: Luciana Elizalde, INBIOMA-CONICET).

### Species identification

Workers were identified based on the knowledge of the current taxonomy, by observing its morphology, using keys (Kusnezov, 1952; Kusnezov, 1978; Snelling & Hunt, 1976), the original species descriptions, comparison with material deposited at the “Colección Entomológica del Instituto-Fundación Miguel Lillo” (IFML) and photographs of type material available (https://www.antweb.org/ and personal of Cuezzo, F.). A stereomicroscope to magnification of no less than 40X was used.

### Library preparation and sequencing

Genomic DNA was extracted for a total of 157 *Dorymyrmex* individuals using the Qiagen DNeasy Blood & Tissue Kit (Qiagen) following the protocol provided by the manufacturer. Purified DNA was quantified using a Qubit 4.0 fluorimeter (Invitrogen, Carlsbad, USA). For each sample, total genomic DNA was digested with the restriction enzymes EcoRI and MseI, adapters and individual barcodes were ligated and samples were pooled in equimolar ratios. The resulting library was purified, size-selected with a range of 290–370 bp using PippinPrep 2% agarose gel cassettes (Sage Science, Sage Science, Beverly, Massachusetts, USA) and amplified by PCR (12 cycles). The final library was sequenced on one lane of a NextSeq500 instrument (Illumina, San Diego, California, USA) using a 150-bp single-end protocol at Fasteris facility (Geneva, Switzerland).

### Genomic data assembly

A total of 240 million (M) raw reads were processed by the process_radtags scripts implemented in the Stacks v2.54 package (Catchen, et al., 2013; Rochette, et al., 2019) to demultiplex individuals while removing low-quality reads (process_radtags options: -c –q). Fifteen individuals were removed on the basis of a pronounced drop in read counts below 240,000. Three additional individuals were removed due to a high proportion of missing SNPs (∼90%), thus the final genomic dataset is composed of 138 individuals (Supplementary Table S1). A *de novo* clustering was performed with dDocent (Puritz, et al., 2014), using type of assembly = PE, clustering similarity 0.9 with minimum individual coverage of 2 (k1 and k2), a mapping match value of 1, a mapping mismatch of 3, and a gap penalty of 5. Complex variants were processed using process_complex.py (Kautt, et al., 2020). Indels were removed and SNPs with a minimum depth 5 (--minDP) were filtered using vcftools v0.1.15 (Danecek, et al., 2011). A final data set of 1.7M reads per individual in average (450k sites and depth 50 in average) was used for downstream analysis (Supplementary Table S1). Missing data was filtered depending on the analysis (as specified below).

### Population statistics, clusters and gene flow analysis

Individual ancestry components were estimated based on 121,703 SNPs with a permissive threshold (--max-missing 0.2) representing all ingroup species (n = 126) was used to retain the largest possible variation, using Admixture v1.3 (Alexander & Lange, 2011) including 5-fold cross-validation (--cv=5). Fixation index F_st_ was calculated using pairwise.pop.fst function of the hierfstat R package (Goudet, 2004) for pairs of sampling points. We estimated inbreeding coefficient using vcftools only from sites with no missing data (--max-missing 1). Windowed nucleotide diversity (π) is more robust to missingness, so a relaxed threshold (--window-pi 150, --max-missing 0.8) was used to retain more sites per window.

### Phylogenetic tree inferences

A neighbor joining tree (NJ) and a coalescent-based phylogenetic tree were inferred using 19,121 SNPs (one random SNP per locus only, maximum missing data 0.5) and including a total of 276 alleles (i.e., 138 diploid individuals). Phylogenetic inferences were performed using the nj command and the coalescent-based program SVDquartets (Chifman & Kubatko, 2014) in PAUP* 4.0 (Swofford, 2003). The analysis was run by sampling one million random quartets and calculating 100 bootstrap replicates to assess statistical support.

### Phylogenetic network inference

A phylogenetic network under the multispecies network coalescent was inferred using the PhyloNetworks program (Solís-Lemus, et al., 2017). The program uses a pseudolikelihood approach and focuses on quartets, calculating the observed quartet concordance factor (CF). The CF of a given quartet (or split) is the proportion trees supporting that particular quartet (Baum, 2007). A random SNP was extracted from each locus and the input file of concordance factors for all possible 4-taxon combinations (330 species quartets in total) was generated using the SNPsCF function (www.github.com/melisaolave/SNPs2CF ; (Olave & Meyer, 2020)). We sampled a total of 50 individual quartets (n.quartets = 50, between.species.only = TRUE) using default values for all remaining settings. The phylogenetic networks were reconstructed using the function snaq! (Solís-Lemus & Ané, 2016) in PhyloNetworks (Solís-Lemus, et al., 2017), including 20 independent runs with random seeds. We used the SVDquartet tree as starting topology, and then inferred the networks using hmax = 0 (no hybridization), 1, 2, 3 and 4. Following the author’s recommendations, each analysis was seeded with the best network from the previous (hmax -1) inference. The best number of hybridization parameters was selected by plotting the likelihood scores and observing the tipping points in the distributions, i.e., a sharp improvement is expected until the number of hybridizing edges reaches the best value and a slower linear improvement thereafter. Finally, concordance factor histograms were plotted to investigate genome wide SNP discordance, using the custom function plotCF of the SNPs2CF package.

### Phylogeographic diversification analysis

To infer diversification across space and time, a continuous phylogeography analysis (Lemey, et al., 2010) was run in BEAST v2.6.7 (Bouckaert, et al., 2019). One individual per species/clade and per sampling locality was selected, leading to a matrix with 28 individuals and their respective geographic coordinates. Individuals with the lowest missing data were preferred. A total of 100 loci were randomly selected based on a previous filtered list of loci obtained using a custom function select.contigs relased in mafalda R package (www.github.com/melisaolave/mafalda), following the criteria: number of SNPs within the 95% confidence interval based on general distribution and <0.1 missing individuals per site in the RAD loci. Loci of 138 bp were reconstructed using our custom function vcf2Loci of mafalda R package. The final matrix consisted of 13,800 bp. Considering a previously proposed mutation rate for ants (Portinha, et al., 2022) based on estimations of haploid social insects’ rates per generation (Liu, et al., 2017), a mutation rate = 2.8 x10^-3^ site/my was used in a strict clock for BEAST analysis, after considering 2.5 years generation time (Pessacq observations) and diploid individuals. A partition with spherical geography was estimated under a relaxed clock with log-normal distribution. The analysis was run with a MCMC of 100 million generations, sampled every 10,000 steps, and 25% burnin. Convergence was assessed with ESS values for parameter estimations equal to or greater than 200. The input xml file is provided as supplementary material.

### Demographic model and species delimitation analysis

A Bayesian coalescent-based demographic model was estimated using bpp v4, including species delimitation analysis (analysis A10; (Flouris, et al., 2020)). Because the three lineage groups were already strongly supported as distinct by the phylogenomic and network analyses and given computational constraints, three separate analyses were run, as follow: (i) *D. cf. richteri* C1 + C2 + *D. cf. tener* C3 + C4; (2) D. *cf. tener* C5 + sp. C7 + *D. cf. tener* C6; and (3) *D. flavescens* C8 + C9 + C10. For each matrix, 200 loci were randomly selected based on a previous filtered list of loci obtained using our custom function select.contigs in mafalda R package, following the criteria: number of SNPs within 95% confidence interval based on general distribution and <0.1 missing individuals per site in the RAD locus. Loci of 138 bp were reconstructed using the custom function vcf2Loci. The SVDquartets topology was used as guide tree, and a diffuse inverse-tau priors IT = (3, 0.015) and a diffuse inverse-gamma prior were adjusted depending on the average observed substitutions per site per individual observed in the 100 loci dataset, as by the authors: matrix (1) with IG = (3, 0.0008), matrix (2) with IG = (3, 0.002) and matrix (3) with IG = (3, 0.004). A rjMCMC was run during 500K generations, with 10% burnin. All remaining settings were used as default. Convergence was assessed on parameter’s ESS > 200. Estimates of theta were converted into effective population sizes (N_e_) based on the above-mentioned mutation rate. The genealogical divergence index (*gdi*, (Jackson, et al., 2017; Leaché, et al., 2019) was inferred using get.gdi function in mafalda R package. The *gdi* is the probability that the two sequences within a population do not coalesce before reaching species divergence (*τ*) when tracing the genealogy backwards in time (Leaché, et al., 2019).

### Genomes scans

The R package PCAdapt (Luu, et al., 2017) was used to calculate a PCA and detect SNPs under selection and candidates of local genomic adaptation. The method performs a PCA to summarize population structure and tests each SNP for association with the retained principal components. Analyses were run for combinations of lineages identified as closely related and divergent in the phylogenetic and species delimitation analyses, allowing 20% missing data and applying a multiple-test Bonferroni correction. RAD loci containing one or more outliers were blasted against NCBI using blastn (max_target_seqs 500, e-value 0.0001) to obtain annotations.

## Results and discussion

### An extensive cryptic diversity

The neighbor joining (NJ) tree displays high genetic structure and, surprisingly, extended paraphyly of species identified as D. *richteri* and D. *tener* based on morphological observations (Fig. 1d), while three clades are detected within D. *flavescens*. The clades were named as originally identified species based on morphology, plus an assignation to clades as from C1 to 10 (Fig. 1). Pairwise F_st_ calculations also show strong differentiation between some clades (ranging from 0.01 to 0.73; Supplementary Table S3; Supplementary Fig. S1). Admixture analysis is in agreement with the NJ tree (Fig. 1b; Supplementary Fig. S1), revealing the existence of seven different groups with different ancestry estimations (K = 7, best supported CV = 0.13464; Supplementary Table S2). However, the analysis still detects recent admixutre between clade pairs (i.e., *D. cf. richteri* C1 and C2, as well as *D. cf. richteri* C2 and *D. cf. tener C3*; plus *D. cf. tener C3* and *D. cf. tener* C4). Individuals inferred with hybrid ancestry also carry higher heterozygosity estimates than expected by chance (Supplementary Fig. 1; Supplementary Table S1; Supplementary Fig. S2). These clades are found both sympatry and parapatry (Fig. 1a). Despite evidence of recent admixture, they remain genetically differentiated, suggesting that independently evolving lineages have emerged even where reproductive isolation is incomplete.

Based on individual-lineage assigments inferred from ancestry analysis in admixture, we inferred a coalescent-based SVDquartets tree (Fig. 2). Paraphyly is recovered in concordance with the NJ tree with high statistical support (bootstrap = 100 for all nodes), suggesting the existance of three independently evolving lineages in *D. cf. richteri* and three in *D. cf. tener*, one *D. sp*. C7, and possibly three in *D. flavescens*; despite the evidence of recent admixture (Fig. 1a). Given that hybridization can mislead phylogenetic inference (Solís-Lemus, et al., 2016; Long & Kubatko, 2018), we inferred the dynamics of hybridization events by reconstructing a phylogenetic network, which accommodates coalenscence and gene flow. This method is more sensitive to older hybridization events, thus helps to get a broader picture of the evolutionary scenario, as a complement to admixture that tends to detect recent hybridization (Pang & Zhang, 2025). The phylogenetic network recovers three hybridizing edges that explains *Dorymyrmex* evolution, involving introgression ranging between 2% to 9% of the genome. Furthermore, *D. cf. tener* C6 and *D. cf. tener* C5 are recovered as closely related taxa although not as sister lineages, a discordant result compared to the coalescent tree demonstrating that historical introgression and incomplete lineage sorting jointly shaped the evolutionary history of these lineages.

**Fig. 2:**
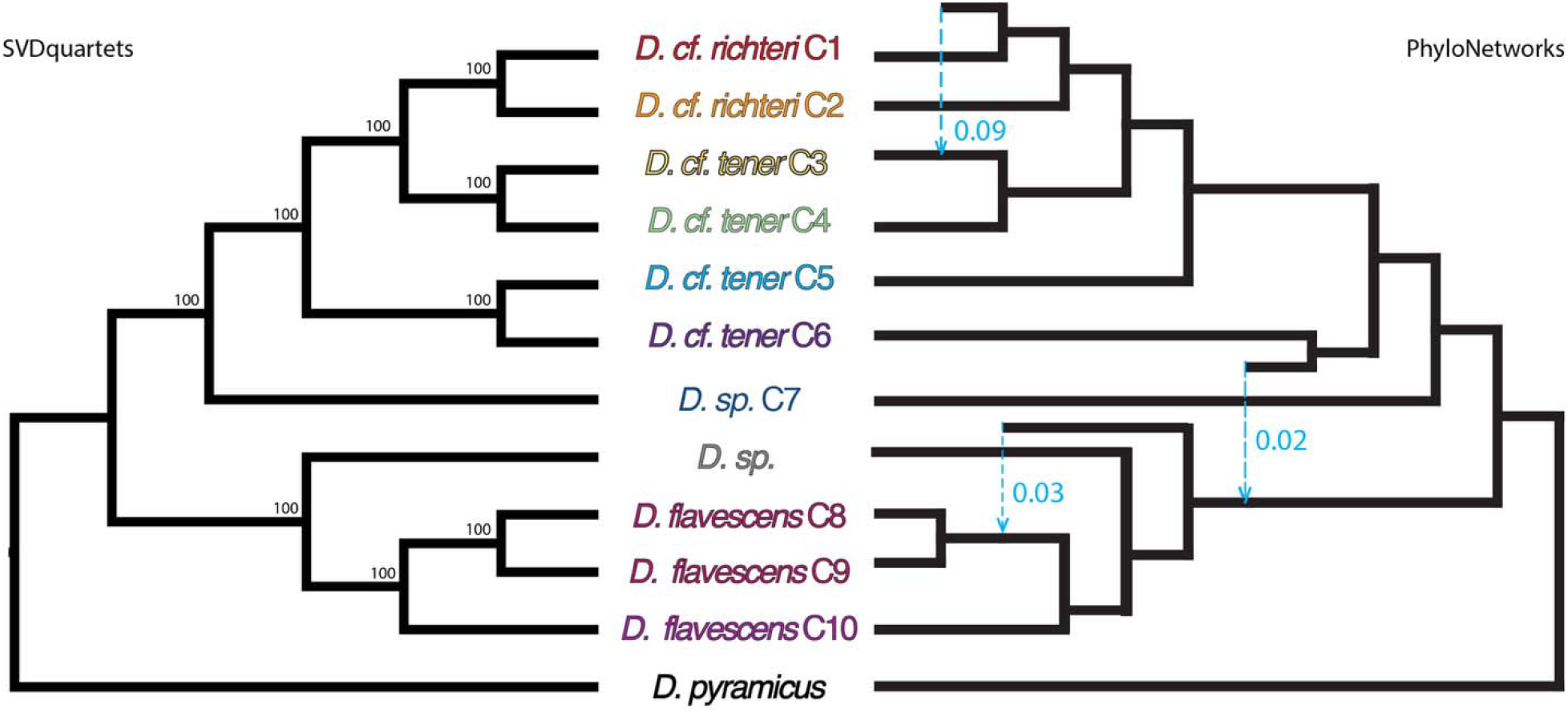
Phylogenomics. SVDquartets coalescent tree (left) and phylogenetic network inference using PhyloNetworks (right), based on 19,121 SNPs for 276 alleles (138 diploid individuals). Bootstrap support is shown on nodes, and inheritance parameter (gamma) is shown in light blue on the phylogenetic network, as the estimated proportion of the genome that has introgressed.

We formally tested species limits, using the coalescent-based program bpp. Results show high posterior probability (>0.95) to support six independent evolving lineages (only the split between the *D. richteri* C1 and C2 is not statistically supported, PP=0.27). In addition, three lineages are detected in *D. flavescens* (C8-C10) with PP=1. As it has been shown that coalescent-based species delimitation models are sensitive to detect population structure (Sukumaran & Knowles, 2017). we calculated the genealogical divergence index (*gdi*, (Jackson, et al., 2017; Leaché, et al., 2019). The *gdi* ranges from 0 (panmixia) to 1 (strong divergence), with values between 0.2 and 0.7 indicating an intermediate stage of divergence, where species boundaries cannot be confidently delimited (the ‘grey zone’ of speciation (Roux, et al., 2016)) usually considered within the gray zone of the speciation continuum, and values above 0.7 are indicative of strong divergence, supporting the existence of “good” species (Jackson, et al., 2017). Even though deciding *what* a species is, is necessarily subjective in current taxonomy (Sukumaran, et al., 2021), *gdi* provides useful and more conservative information on the strength of divergence between two sister clades. Here, results show high *gdi* calculations (generally with *gdi* >0.65) for all pairs of ant lineages displaying PP > 0.95 (Fig. 3; Supplementary Table S4), indicating strong divergence and supporting their recognition as distinct species. Together with the evidence of recent and historical introgression recovered above, these results indicate that substantial genomic divergence has accumulated despite recurrent gene flow. *Diversification in space and time*. The inferred geographic origin of the study group is western Río Negro, followed by lineage expansion both northward and southward into ecologically contrasting regions (Supplementary File S1). During this spatial diversification, lineages likely adapted to different environments. Clades *D. cf. richteri* C1, C2, and C5 reached southern Patagonia, an area characterized by extreme aridity, sparse shrub cover, and extensive bare ground (Soriano, 1956). To the north, the clade *D. cf. tener* C4 reached the Payunia highlands in southwestern Mendoza and northwestern Neuquén (Argentina), a region shaped by intense volcanic and glacial processes (Martinez Carretero, 2004). These contrasting ecological conditions most probably had a role in promoting high levels of endemism among plants, reptiles, and arthropods in the region, aptly described as an “archipelago of mountains” (Martinez Carretero, 2004; Domínguez, et al., 2006; Corbalán & Debandi, 2008; Roig-Juñent, et al., 2008; Olave, et al., 2023; Sánchez, et al., 2026). These distinct ecological landscapes likely imposed strong, contrasting selective pressures, driving adaptive divergence among lineages at the genomic level. Our genome scan analyses comparing closely related lineages and identified between eight and 54 SNPs as outliers, potentially representing signals of local adaptation (Fig. 3). Loci containing these SNPs were extracted and blasted on NCBI, and some of them were found to be annotated genes (Supplementary Table S5). A search was performed on FlyBase, UniProt and NIH to obtain gene functions. Results show that these outlier loci correspond to genes involved in the regulation of cellular and developmental processes, including transcriptional and post-transcriptional control (Rga, FUBP1, EIF4G3, aos, APC4), cell-cycle progression (APC4, RBBP8-like), and signal transduction pathways (CHICO, aos). In addition, some genes are associated with neuronal development and cell architecture, such as axon guidance (Dscam2, ort), cell adhesion (Dscam2), cytoskeletal organization (shot, Nesprin-1), and nuclear–cytoskeletal coupling (Nesprin-1). These functions suggest that genomic divergence is concentrated in regulatory systems controlling gene expression, cellular state, and the integration of environmental signals into developmental and physiological responses. Beyond the specific function of each gene, these candidate loci associated with these heterogeneous environments offer a window into how *Dorymyrmex* lineages colonized such contrasted habitats, consistent with a role for ecological divergence in maintaining genomic differentiation despite recurrent introgression.

**Fig. 3:**
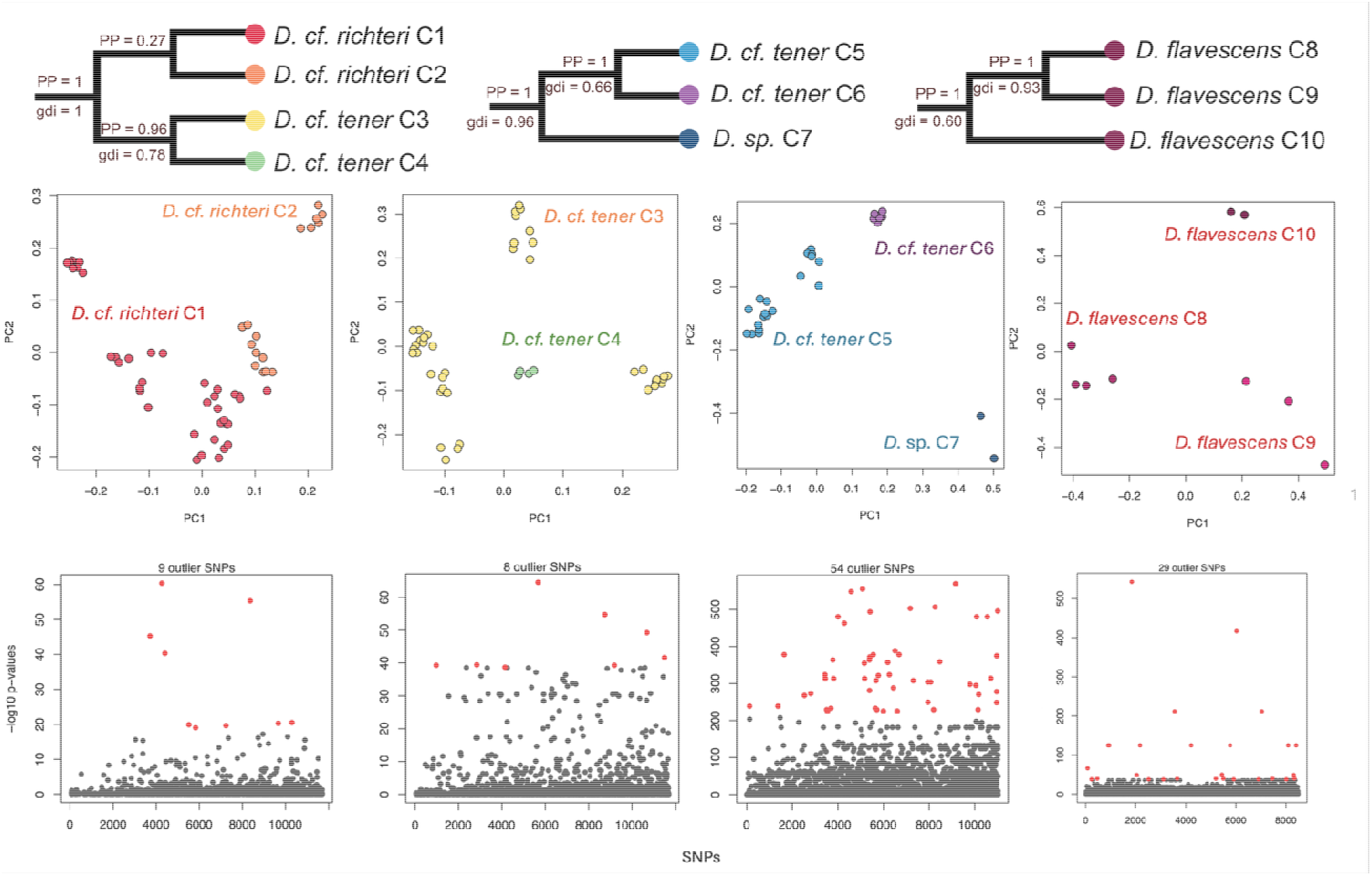
Species delimitation analyses and genome scans. **Top**, result of coalescent-based species delimitation analysis inferred using the bpp program. Posterior probabilities of the split of different lineages are shown on each node as well as *gdi* calculations. Bottom, scatter plots of PCs calculated during genome scans analyzed with PCAdapt, and Manhanttan plots for detections of SNPs outliers, i.e. gene candidates for local adaptations.

### Conservation remarks

Finally, we evaluated demographic parameters to estimate the genetic health of these lineages. Our results indicate that lineages maintain relatively stable estimates of genetic diversity and inbreeding, with no clear genomic signatures of recent inbreeding or critically reduced diversity. This is evidenced by inbreeding coefficients F centered around zero or below (Supplementary Fig. S3) and homogeneous patterns of nucleotide diversity pi (Supplementary Fig. S4), with *D. flavescens* C8–C10 and *D*. sp. C7 carrying the greatest diversity. Even in lineages with comparatively smaller effective population sizes (Supplementary table S6), such as *D. cf. tener* C6, the lack of additional genomic warning signals in genetic diversity and inbreeding, suggests no immediate cause for concern. From a conservation perspective, the overall high heterozygosity and moderate to large effective population sizes are not alarming, as they suggest the maintenance of the adaptive evolutionary potential across lineages. Together with the extensive cryptic diversity uncovered here, these findings emphasize the importance of recognizing previously hidden evolutionary lineages when assessing and conserving Patagonia’s biodiversity.

## Conclusions

The genetic analyses conducted in this study reveal a complex and previously unrecognized evolutionary history within representatives of the Patagonian *Dorymyrmex* ant species. We demonstrate that substantial genomic divergence accumulated despite recurrent historical and contemporary hybridization, resulting in multiple independently evolving lineages, many of which exceed the grey zone of the speciation continuum. *Dorymyrmex* evolution also involves recurrent hybridization appears to have accompanied the diversification of these ants, adding another layer of complexity. The events underscore the challenges of evolutionary inferences posed by genomic discordance due to ancestral polymorphism and hybridization.

Given the insufficiency number of genetic studies on Patagonian insects, our work provides important insights into the region’s genetic diversity and evolutionary dynamics. Although the genomic patterns recovered here indicate that all identified lineages currently maintain good genetic health from a conservation perspective, there is an urgent need for a formal taxonomic revision of *Dorymyrmex*, including the description of the newly identified species. Such efforts are crucial to ensure their recognition within conservation frameworks and to enable the development of effective, lineage-specific management and protection strategies. More broadly, our results show how genome-scale datasets from biodiversity-rich but genomically understudied regions can uncover both hidden evolutionary diversity and the processes shaping it. By revealing that cryptic diversification can proceed despite recurrent gene flow, this study reinforces Patagonia as a valuable natural laboratory for understanding diversification in dynamic landscapes.

## Acknowledgements

We thank all members of the Entomology lab (IADIZA-CONICET) for continuous support. This study was improved thanks to valuable comments made by XX anonymous reviewers. We thanks fauna and protected area authorities from Neuquén, Chubut and Santa Cruz provinces for collection permits.

