## Supplementary Figures for "Cryptic diversification proceeds despite historical and contemporary hybridization in Patagonian ants"

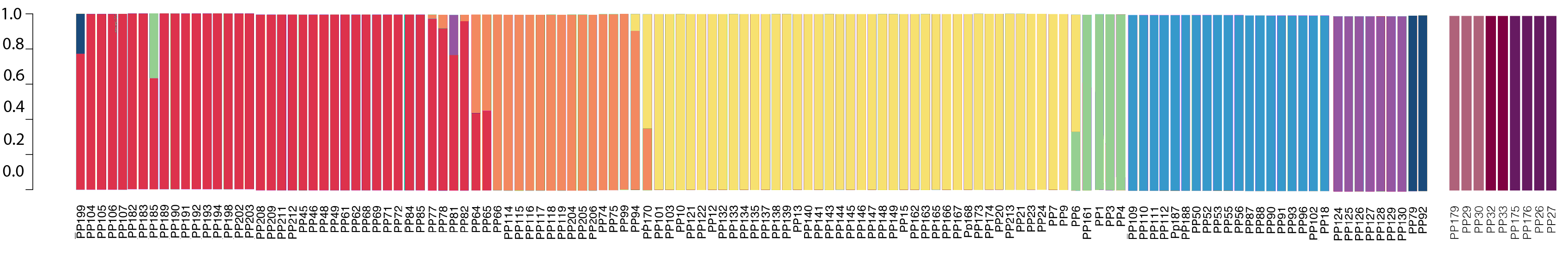

Fig. S1: Full admixture plot constructed on 121,703 SNPs including 126 individuals, including IDs of all samples.

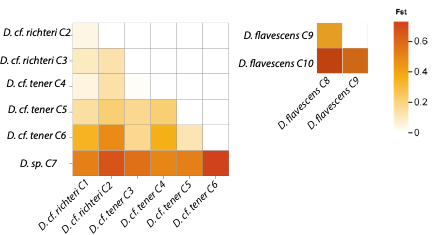

Fig S2: Fst calculations for pairwise comparisions of different lineages recovered.

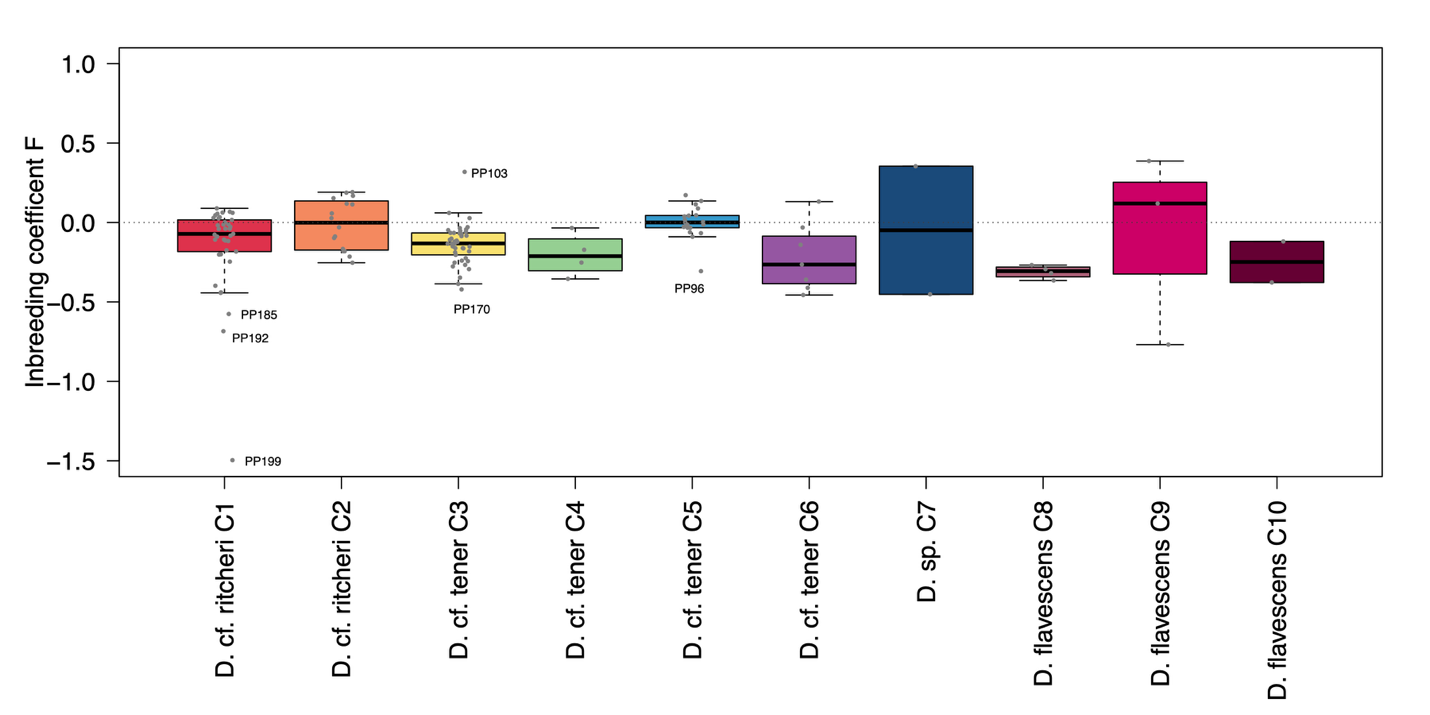

Fig. S3: Inbreeding coefficient F calculated per individual per lineage. Gray dotted line is plotted for reference of F = 0 (equilibrium). Values above zero indicate an excess of homozygosity, and negative values represent an excess of heterozygosity than expected by chance. Sampe IDs for the case of outliers are displayed.

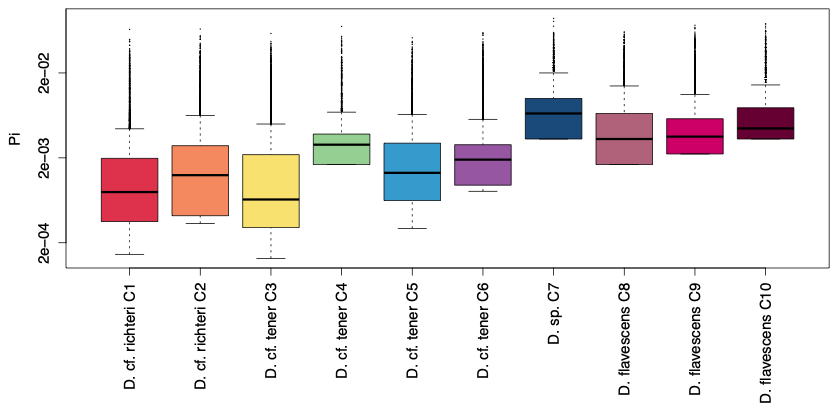

Fig. S4: Genetic diversity (pi) in log scale per lineage estimated per RAD locus. Gray dotted line represents overall mean. Greater values indicate greater genetic diversity.
